# Distinct but cooperating brain networks supporting semantic cognition

**DOI:** 10.1101/2021.07.19.452716

**Authors:** JeYoung Jung, Matthew A. Lambon Ralph

## Abstract

Semantic cognition is a complex brain function involving multiple processes from sensory systems, semantic systems, to domain-general cognitive systems, reflecting its multifaceted nature. However, it remain unclear how these systems cooperate with each other to achieve effective semantic cognition. Here, we investigated the neural networks involved in semantic cognition using independent component analysis (ICA). We used a semantic judgement task and a pattern matching task as a control task with two levels of difficulty to disentangle task-specific networks from domain-general networks and to delineate task-specific involvement of these networks. ICA revealed that semantic processing recruited two task-specific networks (semantic network [SN] and extended semantic network [ESN]) as well as domain general networks including the frontoparietal network (FPN) and default mode network (DMN). Specifically, two distinct semantic networks were differently modulated by task difficulty. The SN was coupled with the extended semantic network and FPN but decoupled with the DMN, whereas the ESN was synchronised with the FPN and DMN. Furthermore, the degree of decoupling between the SN and DMN was associated with semantic performance. Our findings suggest that human higher cognition is achieved by the neural dynamics of brain networks, serving distinct and shared cognitive functions depending on task demands.

## Introduction

Human higher cognition is supported by the dynamic configuration of functional brain networks in response to the environment. Functional brain networks constantly reconfigure their architecture corresponding to cognitive demands in order to achieve successful cognition, which leads to the functional segregation and integration of brain networks (Bressler and Kelso 2001). Importantly, our cognitive performance links to the dynamic reorganization of functional brain networks (Cohen and D’Esposito 2016; Shine et al. 2016). It is one of the important challenges for cognitive neuroscience to understand the dynamics of functional brain networks in relation to cognitive demands. Here, we explored this issue targeting semantic cognition as our cognitive domain of interest given that it is a core higher cognitive function involving complex and multiple cognitive processes.

Semantic cognition refers to our ability to use, manipulate and generalize knowledge in order to interact with the world, by producing time- and context-appropriate behaviours (Binder et al. 2009; Lambon Ralph et al. 2017). It requires a complex brain function involving multiple processes from sensory systems (Binder et al. 2016; Martin 2016), semantic representation (Binney et al. 2010; Lambon Ralph, Sage, et al. 2010; Chen et al. 2016), controlled semantic retrievals (Whitney et al. 2011; Noonan et al. 2013), to domain-general cognitive systems (Raichle et al. 2001; Duncan 2010; Vossel et al. 2014). In addition to task-active systems, the default mode network (DMN), a task negative system, is also engaged in accessing episodic and semantic memories, without goal-directed semantic retrieval (Binder et al. 1999). Although these distinctive systems are involved in semantic processing, it remain unclear how these systems cooperate with each other to achieve effective semantic cognition.

Lambon Ralph and colleagues (2017) proposed a framework called ‘controlled semantic cognition’ (CSC) to account for two key systems of semantic cognition: semantic representation and control system. In CSC, semantic representation is supported by a distributed system consisting of a transmodal hub and modality-specific spokes. Empirical and computational evidence has demonstrated that the bilateral anterior temporal lobes (ATLs) are a site for the hub of semantic representation (Coccia et al. 2004; Binney *et al*. 2010; Peelen and Caramazza 2012). Damage to this region incurs a severe impairment in semantic knowledge across all modalities (Bozeat et al. 2000; Patterson et al. 2007). Investigations of the structural connectivity in the temporal lobe have demonstrated the convergence of major white matter pathways in the ATL, terminating different parts of the ATL (Binney et al. 2012; Bajada et al. 2017; Jung et al. 2017), supporting the graded ATL-hub proposal (Rice, Hoffman, et al. 2015; Lambon Ralph *et al*. 2017). For example, the temporopolar cortex is connected to the orbitofrontal cortex (OFC) and pars orbitalis via the uncinated fasciculus; superior ATL regions are connected to other prefrontal cortex through the extreme capsule complex and the inferior parietal lobe (IPL) via the middle longitudinal fasciculus; the ventral ATL is linked to the IFC through the inferior longitudinal fasciculus. Local U-fibre connections within the ATL provide the basis for convergence of information from different white-matter pathways, which is maximal in the lateral and ventral ATL regions. Also, resting-state and task-active fMRI studies have reported the similar patterns of ATL connectivity (Pascual et al. 2015; Jackson et al. 2016). Semantic control system guides the representational system to select a particular concept or generate a proper behaviour in a given task or context. For example, writing an email with a keyboard requires fine-motor coordination with hands, *typing*. But if you want to move the keyboard, you have to ignore the main function, *typing*, and produce a different set of hand and arm movement, *grasping and carrying*. The control system is implemented in a distributed system including the left inferior frontal gyrus (IFG), posterior middle temporal gyrus (pMTG) and IPL (Thompson-Schill et al. 1997; Badre and Wagner 2002; Noonan *et al*. 2013). Damage to this system observed in semantic aphasia (SA) produces difficulty controlling conceptual retrieval to suit a task or context (Jefferies and Lambon Ralph 2006; Corbett et al. 2009).

The semantic control system overlap with the frontoparietal network (FPN) involved in cognitive control across domains (Duncan 2010; Fedorenko et al. 2012). The FPN responds to a wide range of demanding cognitive tasks, which includes the bilateral dorsolateral prefrontal cortex (DLPFC), inferior frontal sulcus (IFS), anterior cingulate cortex (ACC), pre/supplementary motor area (preSMA/SMA), and intraparietal sulcus (IPS). Possibly, the processes involved in semantic control may overlap with domain-general executive control processing. Researchers attempted to delineate these two systems and suggested a superior-inferior functional specialization of the FPN. fMRI studies have reported that the ventral PFC (vPFC) and pMTG showed increased activation during the retrieval of weak semantic association, whereas the DLPFC and IPS were activated for the selection demands regardless of tasks (semantic and non-semantic tasks) (Badre et al. 2005; Nagel et al. 2008). Studies of structural and functional connectivity also showed that the vPFC and pMTG were connected to the ATL, suggesting their role of the regulation of semantic representation, whereas superior parts of the network were not (Jackson *et al*. 2016; Jung *et al*. 2017). However, the relationship between domain-specific and domain-general networks and the spatial overlap of these networks still remain unclear.

Previous fMRI studies have showed that the semantic systems overlap with the DMN (Binder *et al*. 1999; Binder *et al*. 2009; Wirth et al. 2011; Humphreys et al. 2015; Jackson *et al*. 2016). The DMN is localised primarily to midline anterior and posterior cortical regions, angular gyrus (AG), and medial and lateral temporal cortices (Raichle *et al*. 2001). It normally shows activation at rest and deactivation during goal-directed tasks and is involved in autobiographical memory retrieval, self-related thinking, and consciousness (Buckner et al. 2008). Thus, the rich conceptual processing at rest requires the semantic system, showing common brain regions for these two networks (Binder *et al*. 1999). A recent fMRI study investigated the DMN in relation to semantic processing (Wirth *et al*. 2011). The DMN showed less deactivation during semantic processing compared to non-semantic processing, specifically within the left-hemispheric DMN regions including the ATL, AG, medial PFC (mPFC), pMTG, and retrosplenial cortex. Another fMRI study demonstrated different functions of the overlapping regions between the DMN and semantic network (SN) (Humphreys *et al*. 2015). The ATL was activated for all semantic tasks but deactivated for non-semantic tasks, whereas the AG was deactivated for all tasks, showing differential level of deactivation related to task difficulty. These studies suggest the functional segregation and integration of the DMN during semantic processing but it is still poorly understood how these two systems interact with each other to achieve efficient semantic cognition.

Independent component analysis (ICA) is a signal processing technique to extract signals by unmixing signal mixtures (McKeown and Sejnowski 1998; Calhoun, Adali, Pearlson, et al. 2001). ICA assumes that fMRI signal from each voxel represents a linear mixture of source. Then, it separates this signal mixture into independent source signal and groups all voxels into independent components (ICs), which represent temporally coherent functional brain networks. Several studies demonstrated that the ICA revealed more brain regions involved in tasks compared to traditional general linear model (GLM) analysis and different task-related modulation in overlapping regions of two or more functional networks (Calhoun, Adali, McGinty, et al. 2001; Kim et al. 2009; Domagalik et al. 2012; Geranmayeh et al. 2014). Geranmayeh and colleagues (2014) identified overlapping networks during spoken language production and specified their involvement in speech processing, using ICA. Therefore, this approach can be more sensitive to detect task-related brain activity hidden from the GLM approach and delineate specific involvement of overlapping regions according to tasks.

Here, we performed an fMRI study to test for the presence of functionally independent but spatially overlapping brain networks for semantic cognition. We used a semantic judgement task and a pattern matching task as a control task with two different levels of difficulty in order to disentangle domain-specific (semantic) processing from domain-general processing (non-semantic) and to delineate task-specific involvement of overlapping brain networks during semantic processing. We hypothesized that ICA would identify several networks related to semantic processing including the semantic system, FPN, and DMN and task difficulty modulation would segregate or integrate these networks. In order to investigate the functional interaction between these networks during semantic processing, we performed functional network connectivity analysis. Finally, we explored the relationship between the internetwork connectivity and semantic task performance.

## Methods

### Participants

Twenty three healthy young participants were recruited for this study (12 females, mean age = 21 ± 3 years ranging from 19 to 30 years). They were English native speakers with right-handedness (Oldfield 1971) and had normal or corrected-normal vision. All participants gave their written informed consent prior to the study. The experiment was approved by local ethics committee in accordance with the Declaration of Helsinki.

### Experimental design and procedure

Participants performed a semantic judgement task and a pattern matching task (a control task) with two levels of difficulty (easy vs. hard) during fMRI. The semantic judgement task was adapted from a previous fMRI study (Jackson et al. 2015). Participants were presented with triads of concrete nouns and asked to judge which of the two choices was more related to the probe word. In each trial, 3 words were presented on the screen, a probe on the top and 2 choices (a target and a foil) at the bottom. The level of difficulty was modulated by two different foils. One foil was from an unrelated category to the probe word (easy condition) (e.g., CARROT [probe] – GRAPE [target] paired with the foil, TELESCOPE). The other foil was from the same or a related category to the target and probe (hard condition) (e.g., CARROT [probe] – GRAPE [target] paired with the foil, BLUBELL). Targets, probes and foils were matched for frequency, imagebility, letter length, and syllable length (ps > 0.5). A pattern matching task was also adapted from a previous study (Jung et al. 2021). The items for the control task were generated by visually scrambling items from the semantic task. Each pattern was created by scrambling each item into 120 pieces and re-arranging them in a random order. Participants were asked to select which of 2 patterns was identical to a probe pattern. In the hard condition, the target patterns were presented 180° rotated.

In the scanner, stimuli were presented in a block design and each block contained 4 trials from each condition (semantic easy [SE], semantic hard [SH], control easy [CE] and control hard [CH]). Between the task blocks, there were fixation blocks for 4000ms. Each trial started with a 500ms fixation and was presented for 3000ms. Participants responded by pressing one of 2 buttons representing the left and right options. The four conditions were sampled 7 times, giving a total of 28 blocks. The semantic and control tasks were paired for the same difficulty level and presented in a counterbalanced order (e.g., A-B-A-B). The total time of scanning was about 8mins. E-prime software (Psychology Software Tools Inc., Pittsburgh, USA) was used to display stimuli and to record responses.

### fMRI data acquisition and general linear model (GLM) analysis

Imaging was performed on a 3T Philips Achieva scanner using a 32-channel head coil with a SENSE factor 2.5. To maximise signal-to-noise (SNR) in the ATL, we utilised a dual-echo fMRI protocol developed by Halai et al (Halai et al. 2014). The fMRI sequence included 42 slices, 96 × 96 matrix, 240 × 240 × 126mm FOV, in-plane resolution 2.5 × 2.5, slice thickness 3mm, TR = 2.8s, TE = 12ms and 35ms. 180 dynamic scans were acquired. The structural image was acquired using a 3D MPRAGE pulse sequence with 200 slices, in planed resolution 0.94 × 0.94, slice thinkness 0.9mm, TR = 8.4ms, and TE = 3.9ms.

Analysis was carried out using SPM8 (Wellcome Department of Imaging Neuroscience, London; www.fil.ion.ucl.ac.uk/spm). The dual gradient echo images were extracted and averaged using in-house MATLAB code developed by Halai et al (Halai *et al*. 2014). Functional images were realigned correcting for motion artefacts and different signal acquisition times by shifting the signal measured in each slice relative to the acquisition of the middle slice prior to combining the short and long echo images. The mean functional EPI image was co-registered to the individual T1-weighted image and segmented using the DARTEL (diffeomorphic anatomical registration through an exponentiated lie algebra) toolbox (Ashburner 2007). Then, normalization was performed using DARTEL to warp and reslice images into MNI space and smoothing was applied with an 8mm full-width half-maximum Gaussian filter.

At the individual subject level, contrasts of interest were modelled using a box-car function convolved with the canonical hemodynamic response function. Four separate regressors were modelled according to task and difficulty (SE, SH, CE and CH). At the group level, a two-factorial ANOVA with task (semantic vs. control) and difficulty (easy vs. hard) was conducted for the main effect of task and interaction between task and difficulty and T-contrasts were established for the contrast of semantic > control and control > semantic. Whole-brain maps were thresholded at p < 0.001 at the voxel level, with a FWE-corrected cluster threshold of p < 0.05, ks > 100.

### Independent component analysis (ICA)

ICA is a data-driven multivariate approach to decompose a mixed signal into independent components (ICs) (Calhoun, Adali, Pearlson, *et al*. 2001). ICA utilizes fluctuation in the fMRI data to separate the signal into maximally independent spatial maps (components), each explaining unique variance of the 4D fMRI data. Each component has a time course related to a coherent neural signalling associated with a specific task, artifact, or both.

ICA was performed using a group ICA algorithm (GIFT, http://icatb.sourceforge.net/, version 3.0a). Using Maximum Description Length (MDL) and Akaike’s criteria, the number of independent components was estimated. A first stage subject-specific principal components analysis (PCA) was performed. A second stage group data reduction, using the expectation-maximization algorithm included in GIFT. Then, Informax ICA algorithm (Bell and Sejnowski 1995) was conducted, repeating it 20 times in ICASSO implemented in GIFT to generate a stable set of 30 final components. Finally, the ICs were then estimated using the GICA back-reconstruction method based on PCA compression and projection (Calhoun, Adali, Pearlson, *et al*. 2001).

Of the resultant 30 ICs, 15 components were related to residual artifact including the signal distributed around the edge of the brain and within cerebrospinal fluid spaces, variation in head size, or vascular blood flow. These were excluded for further analysis. The rest 15 ICs had the signal distributed within the brain and were defined as ‘networks’. We labelled them with regional or functional descriptors (e.g., default mode network, frontoparietal network).

For each of the 15 components, we tested whether that component was significantly functionally involved in any task condition, using the ‘temporal sort’ of GIFT. Temporal sorting was conducted by applying a GLM to the component’s time course. The fMRI run specific time course for each subject were regressed against the design matrix for the tasks and tested for significance to identify components where activity was greater during semantic and control processing (easy and hard) against rest. The resulting beta (β) weights represent the degree to which component network recruitment was modulated by the task conditions. For a given component, positive and negative β weights indicate task-related network recruitment that is increased or decreased with respect to baseline, respectively. To evaluate the task-related network recruitment, we performed a one-sample t-test on the β weights (p _FDR-corrected_ < 0.005). A two-factorial ANOVA with task (semantic vs. control) and difficulty (easy vs. hard) was conducted for the main effect of task and interaction between task and difficulty. For the comparison between easy and hard condition for each task, *post hoc* paired t-tests were performed for each component (p < 0.05).

We evaluated the spatial similarity between the components and GLM brain activation maps, using the ‘spatial sort’ of GIFT. Spatial sorting was conducted on the components showed the significant task-relatedness in temporal sorting, by applying the thresholded GLM group results from the contrasts of interest to the component’s spatial maps. The resulting beta (β) weights represent the degree to which component network was overlapping with the GLM brain activation maps. To evaluate the spatial similarity, we performed a one-sample t-test on the β weights (p _FDR-corrected_ < 0.005). For the comparison between easy and hard condition for each task, paired t-tests were performed for each component (p _FDR-corrected_ < 0.05).

To assess the connectivity between networks, functional network connectivity (FNC) was performed for the networks showed the significant temporal and spatial task modulations. FNC was estimated as the Pearson’s correlation coefficients between pairs of time-courses of networks (Jafri et al. 2008) and tested using one-sample t-test (p _FDR-corrected_ < 0.05). Then, to explore the relationship between the FNC and semantic performance, correlation analysis was performed (p _FDR-corrected_ < 0.05).

## Results

### Behavioural results

A repeated-measure ANOVA with task (semantic vs. control) and difficulty (easy vs. hard) was conducted for accuracy and reaction time (RT). In accuracy, there was a significant main effect of difficulty (F_1, 22_ = 18.34, p < 0.001). The other main effect and interaction did not reach the significance. In RT, the results showed a significant main effect of task (F_1, 22_ = 11.54, p < 0.005), difficulty (F_1, 22_ = 64.22, p < 0.001) and an interaction (F_1, 22_ = 24.67, p < 0.001). *Post hoc* paired t-tests revealed that the difficulty manipulation was successful in the accuracy and RT. The accuracy was significantly reduced (semantic: t = 6.23, p < 0.001; control: t = 2.35, p < 0.05) and the RT was significantly increased for the hard condition compared to the easy condition (semantic: t = −4.80, p < 0.001; control: t = -8.52, p < 0.001). There was no difference in the accuracy between semantic and control tasks (ps > 0.35). The RT showed no difference in the easy condition (p = 0.69) but a significant difference in the hard condition between the semantic and control tasks (t = -5.56, p < 0.001). The results are summarised in Table 1.

**Table 1.** Behavioural results. mean ± standard error

|  | Semantic |  | Control |  |
| --- | --- | --- | --- | --- |
|  | ACC | RT | ACC | RT |
| Easy | 93.2 ± 1.6 | 1411.2 ± 51.3 | 90.8 ± 3.1 | 1430.7 ± 48.9 |
| Hard | 83.9 ± 1.8 | 1603.4 ± 45.2 | 81.3 ± 2.3 | 1859.1 ± 40.9 |

### Imaging results

The ICA revealed 15 components showing the patterns of temporally coherent signal within the brain. We refer to these components as brain networks and label them according to their spatial location (e.g., frontoparietal network: FPN) or previously described labels (e.g., default mode network: DMN). Figure 1 and Table S1 summarise the results of these 15 components.

**Figure 1.**
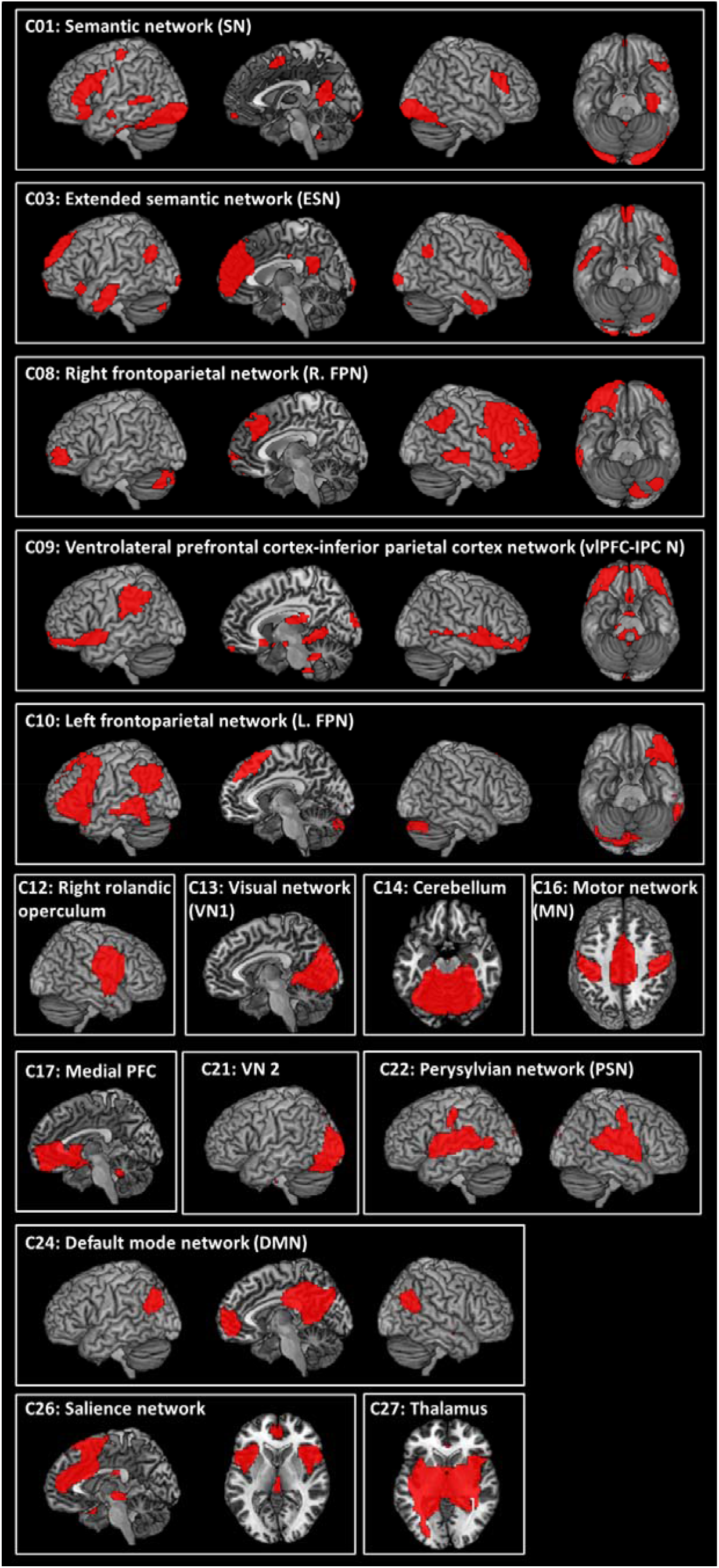
The ICA results. Statistical threshold was set at z > 4. See Table S1 for coordinates.

Figure 2 and Table 2 display the ICA results of temporal regression analysis, showing how the different networks are modulated by the task conditions. Semantic conditions (easy and hard) significantly activated C01 (semantic network: SN), C03 (extended semantic network: ESN), C10 (L.FPN), and C22 (Perysylvian network: PSN) and suppressed C13 (visual network 1: VN1), C21 (VN2), and C24 (DMN). C27 (thalamus) was significantly activated by the hard semantic condition only. In contrast, both control conditions significantly activated C13 and C21 and suppressed the networks activated by semantic processing (C01, C06, C10, and C22). The easy control condition inhibited C12 (rolandic operculum), whereas the hard condition activated C26 (salience network) and supressed C27. The ICA results of spatial regression analysis is summarised in Table S2.

**Figure 2.**
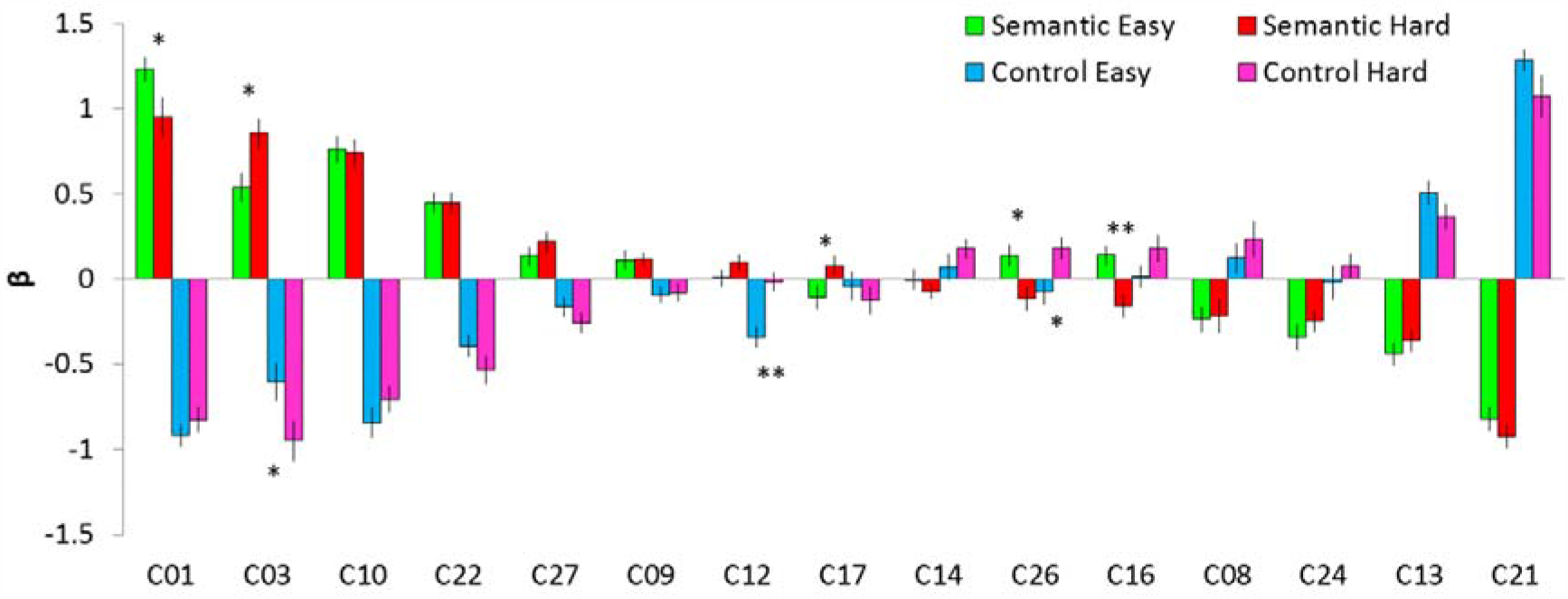
The results of temporal regression analysis. Bar graph shows the mean β weight for each condition against the baseline. Green and red bars represent semantic easy and hard conditions, respectively. Cyan and pink bars represent control easy and hard conditions, respectively. Error bars indicate standard errors. Significant difference between easy and hard conditions is shown with * p < 0.05, ** p < 0.01.

**Table 2.** The result of ICA temporal regression. Bold indicates the significant results from one-sample t-tests. P _FDR-corrected_ < 0.005

| IC | Semantic Easy |  | Semantic Hard |  | Control Easy |  | Control Hard |  |
| --- | --- | --- | --- | --- | --- | --- | --- | --- |
|  | T-value | p | T-value | p | T-value | p | T-value | p |
| C01 | **17.06** | **0.0000** | **8.07** | **0.0000** | **-15.15** | **0.0000** | **-10.95** | **0.0000** |
| C03 | **6.78** | **0.0000** | **8.78** | **0.0000** | **-5.98** | **0.0000** | **-7.12** | **0.0000** |
| C08 | -3.17 | 0.0044 | -2.19 | 0.0393 | 1.53 | 0.1397 | 2.06 | 0.0515 |
| C09 | 2.02 | 0.0554 | 3.11 | 0.0051 | -1.89 | 0.0726 | -1.60 | 0.1230 |
| C10 | **10.21** | **0.0000** | **9.37** | **0.0000** | **-9.88** | **0.0000** | **-9.10** | **0.0000** |
| C12 | 0.05 | 0.9620 | 2.11 | 0.0469 | **-5.84** | **0.0000** | -0.36 | 0.7209 |
| C13 | **-6.58** | **0.0000** | **-5.59** | **0.0000** | **7.18** | **0.0000** | **5.06** | **0.0000** |
| C14 | -0.02 | 0.9816 | -1.79 | 0.0871 | 0.96 | 0.3470 | 2.89 | 0.0086 |
| C16 | 2.56 | 0.0177 | -2.32 | 0.0302 | 0.20 | 0.8449 | 2.41 | 0.0246 |
| C17 | -1.47 | 0.1570 | 1.35 | 0.1903 | -0.51 | 0.6151 | -1.57 | 0.1297 |
| C21 | **-11.71** | **0.0000** | **-14.15** | **0.0000** | **19.46** | **0.0000** | **9.03** | **0.0000** |
| C22 | **7.88** | **0.0000** | **7.99** | **0.0000** | **-6.20** | **0.0000** | **-6.56** | **0.0000** |
| C24 | **-4.46** | **0.0002** | **-4.15** | **0.0004** | -0.19 | 0.8485 | 1.12 | 0.2751 |
| C26 | 2.22 | 0.0371 | -1.67 | 0.1084 | -1.03 | 0.3153 | **3.34** | **0.0029** |
| C27 | 2.50 | 0.0205 | **3.53** | **0.0019** | -2.89 | 0.0086 | **-4.09** | **0.0005** |

### Networks related to semantic processing

Figure 3 shows the brain activation maps from the GLM analysis and ICs related to semantic processing. The GLM revealed that the easy semantic condition evoked significant activation in the inferior frontal gyrus (IFG), ventrolateral anterior temporal lobe (vATL), posterior middle temporal gyrus (pMTG), fusiform gyrus (FG) and hippocampus in the left hemisphere as well as the inferior parietal lobe (IPL) and rolandic operculum (RO) in the right hemisphere. The hard semantic condition induced more widespread activation in the same regions found in the easy semantic condition and additional activation in the bilateral angular gyrus (AG), the right vATL, the medial prefrontal gyrus (mPFC), the middle cingulate cortex (MCC), and the pre/postcentral gyrus (Fig 3A and Table S3). In order to evaluate the difficulty effect in the key semantic regions, regions of interest (ROI) analysis was performed in the vATL, IFG, and pMTG. The results showed that the hard semantic condition increased the activation in the left IFG and the right vATL significantly compared to the easy condition (p < 0.05). Also, there was a marginally significant increased activation in the left vATL and right pMTG during the hard semantic processing (p = 0.07) (Fig S1).

**Figure 3.**
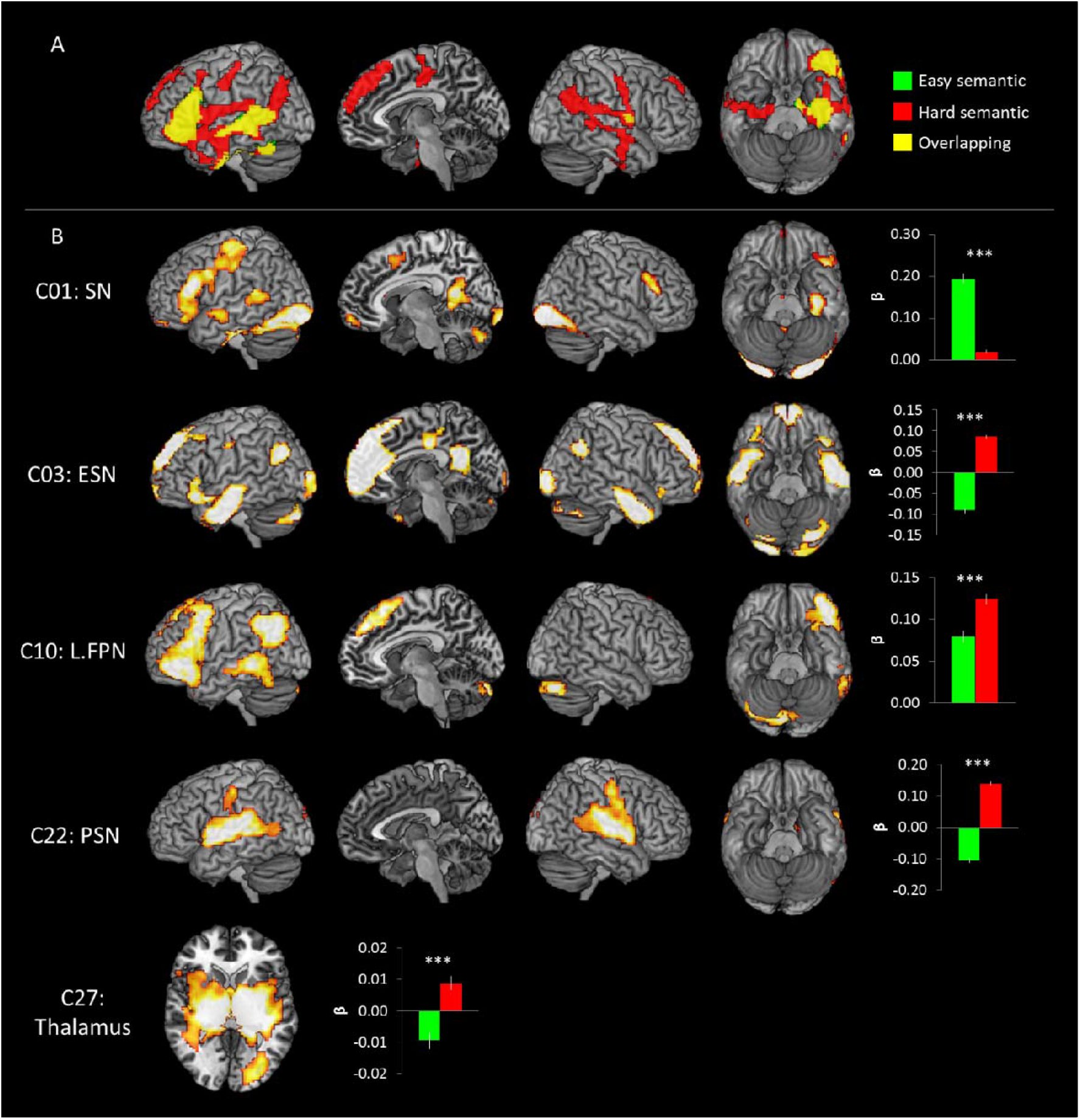
A) The GLM results of the semantic task. Green colour indicates brain regions activated during the easy semantic processing and red colour indicates brain areas during the hard semantic processing. Yellow colour represents the overlapping regions. B) The ICA results of the semantic task. The temporal regression analysis identified that C01, C03, C10, C22, and C27 were significantly involved in the semantic task. The spatial regression analysis showed the spatial overlapping between the GLM results and the ICs. Green bar represent the spatial β weights for the easy condition and red bar for the hard condition. *** p _FDR- corrected_ < 0.001

Different from the results of the GLM analysis, ICA revealed that 5 independent networks were involved in semantic processing, differently modulated by the task difficulty (Fig 2 and Fig 3).

C01 (SN) showed the biggest activation in semantic processing and deactivation in control processing. Similar to the GLM results, C01 consisted of the key semantic regions including the left vATL, left IFG (p. triangularis p. orbitalis, and p. opercularis), left pMTG, mPFC, SMA, bilateral FG, and precuneus. The temporal β weights of the network were analysed by 2 × 2 ANOVA with task (semantic vs. control) and difficulty (easy vs. hard) and the results showed a significant main effect of task (F_1, 22_ = 216.01, p < 0.001), difficulty (F_1, 22_ = 6.59, p < 0.05) and an interaction (F_1, 22_ = 4.73, p < 0.05). *Post hoc* paired t-tests revealed that C01 was significantly more involved in the easy semantic condition compared to the hard condition (p < 0.05) (Fig 2). Consistent to the results of temporal regression, spatial regression analysis demonstrated that the spatial overlapping was significantly greater between C01 and the easy semantic GLM result than the hard semantic result (Fig 3B).

In contrast to C01, C03 (ESN) showed greater temporal and spatial engagement in the hard semantic condition compared to the easy condition. C03 included the bilateral ATL, left IFG (p. triangularis and p. orbitalis), AG, mPFC, superior frontal gyrus (SFG), anterior/middle/posterior cingulate cortex (ACC/MCC/PCC), and visual cortex. 2 × 2 ANOVA with task and difficulty on the temporal β weights demonstrated a significant main effect of task (F_1, 22_ = 141.66, p < 0.001) and an interaction (F_1, 22_ = 4.87, p < 0.05). *Post hoc* paired t-tests revealed that C03 was significantly greater involvement in the hard semantic condition (p < 0.05) (Fig 2). Also, the spatial β weights were negative during the easy processing and positive during the hard processing. It indicates that C03 was more involved in the hard semantic condition, showing greater overlapping between the spatial maps (IC and GLM result derived from the hard processing) (Fig 3).

C10 (L.FPN) includes the left lateral prefrontal cortex, left IPL, and left pMTG and was activated for semantic processing, regardless of task difficulty. L. FPN is active for a wide range of tasks and thought to be involved in modulating cognitive control (Zanto and Gazzaley 2013). However, we found that C10 was activated by the semantic task and suppressed by the control task (Fig 2). The temporal β weights ANOVA revealed a significant main effect of task (F_1, 22_ = 219.23, p < 0.001) and difficulty (F_1, 22_ = 5.06, p < 0.05). Contrary to the temporal regression, the spatial regression showed that C10 was overlapped with the brain activation map of the hard semantic condition more than that of the easy semantic condition (Fig 3B).

C22 (PSN) showed significant semantic-related activity regardless of task difficulty (Fig 2). The PSN includes the bilateral superior temporal gyrus, insular, RO, and the right supramarginal gyrus (SMG). The temporal β weights ANOVA showed a significant main effect of task (F_1, 22_ = 121.27, p < 0.001) and difficulty (F_1, 22_ = 8.23, p < 0.01). Although the temporal regression did not show a significant difference between easy and hard semantic condition, the spatial regression demonstrated that C22 was significantly overlapped with the GLM brain activation map from the hard semantic processing but not with the GLM result of the easy processing (Fig 3B).

C27 (thalamus) consists of the thalamus, putamen, and pallidum. One-sample t-test revealed that the thalamus was significantly involved in the hard semantic condition (Fig 2). The temporal β weights ANOVA showed a significant main effect of task (F_1, 22_ = 25.21, p < 0.001). The spatial β weights analysis showed that C27 was negatively associated with the easy semantic condition, whereas positively related with the hard semantic processing (Fig 3B).

A number of other components either failed to activate for semantic processing or showed deactivations (Fig 2). C17 (mPFC) showed a significant differential involvement in semantic processing according to the task difficulty. C17 was inactive for the easy semantic condition, whereas active for the hard condition. In contrary, C16 (motor network: MN) and C26 (salience network) showed the opposite pattern; active for the easy condition and inactive for the hard condition.

### Networks related to visuo-spatial processing

Figure 4 shows the brain activation maps from the GLM analysis and ICs related to visuo-spatial processing (pattern matching). The GLM demonstrated that the easy control condition induced significant activation in the superior frontal gyrus (SFG), precuneus, and the bilateral visual cortices including the superior/middle occipital gyrus, FG, calcarine gyrus, and lingual gyrus. The hard condition also evoked the activation in the same regions found in the easy condition and in the middle frontal gyrus (MFG), insular, right IFG and right middle orbital gyrus (Fig 4A and Table S3).

**Figure 4.**
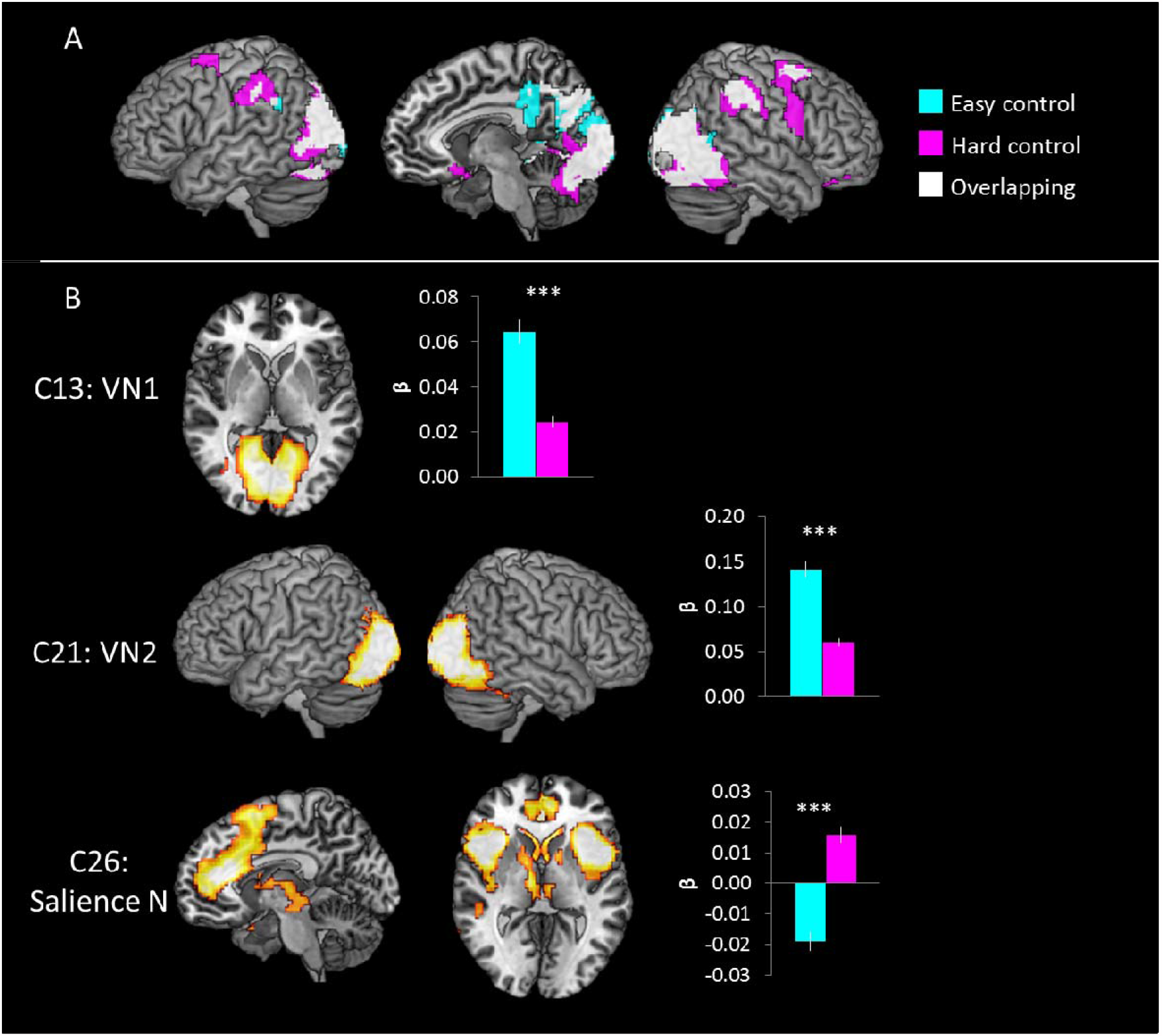
A) The GLM results of the control task. Cyan colour indicates brain regions activated during the easy control processing and pink colour indicates brain areas during the hard control processing. White colour represents the overlapping regions. B) The ICA results of the control task. The temporal regression analysis identified that C13, C21 and C26 were significantly involved in the control task. The spatial regression analysis showed the spatial overlapping between the GLM results and the ICs. Cyan bar represent the spatial β weights for the easy condition and pink bar for the hard condition. *** p _FDR- corrected_ < 0.001

ICA revealed that 3 independent networks were involved in pattern matching processing and were differently modulated by the task difficulty (Fig 2 and Fig 4B). C13 (VN1) including the calcarine gyrus and lingual gyrus was significantly activated by the control task. 2 × 2 ANOVA with task and difficulty on the temporal β weights demonstrated a significant main effect of task (F_1, 22_ = 64.86, p < 0.001) only. The spatial regression showed that C13 was significantly overlapping with the brain activation map of the easy condition than the hard condition (Fig 4B). Similar to C13, C21 (VN2) was significantly associated with the pattern matching processing during the easy condition. C21 consisting of the middle/inferior occipital gyrus and FG showed a significant main effect of task (F_1, 22_ = 327.33, p < 0.001) and difficulty (F_1, 22_ = 16.54, p < 0.001) in the temporal regression analysis (Fig 2). There was no significant effect of interaction. The spatial regression revealed that C21 also was involved in the easy condition more than the hard condition (Fig 4B). In contrast to the visual networks, C26 (salience network) showed significant control task relate activity only for the hard condition (Fig 2). C26 includes the ACC, SMA, middle frontal gyrus, insular, and caudate nucleus and is thought to be critical for detecting behaviourally relevant stimuli and for coordinating the brain’s neural resources in response to these stimuli (Uddin 2017). The temporal β weights of C26 revealed a significant interaction effect between task and difficulty (F_1, 22_ = 8.59, p < 0.01). *Post hoc* paired t-tests revealed that C26 was significantly greater involvement in the hard control condition (Fig 2). The spatial β weights analysis showed that C26 was negatively associated with the easy control condition, whereas positively related with the hard visual processing (Fig 4B).

### Networks related to the interaction between task and difficulty

The interaction effect between task and difficulty evoked the significant activation in the left IFG, left AG, medial prefrontal cortex (PFC), precuneus, right inferior parietal sulcus (IPS), right middle occipital gyrus (MOG), and right inferior temporal gyrus (ITG) (Fig 5A and Table S1). Specifically, the left IFG and ventral/dorsal mPFC showed the increased activation only when the semantic task was hard. Hard semantic processing reduced the deactivation in the AG and precuneus compared to the easy processing (Fig 5A). These regions were more deactivated during the hard control processing. The right IPS and MOG were activated by the control task and more activated during the hard control condition. Overall, the left and medial parts of brain region showed a preference for the semantic processing and demanding semantic processing reduced the deactivation and increased activation in these regions. In contrary, the right hemispheric regions favours for visual processing and difficulty manipulation increased their regional activity.

**Figure 5.**
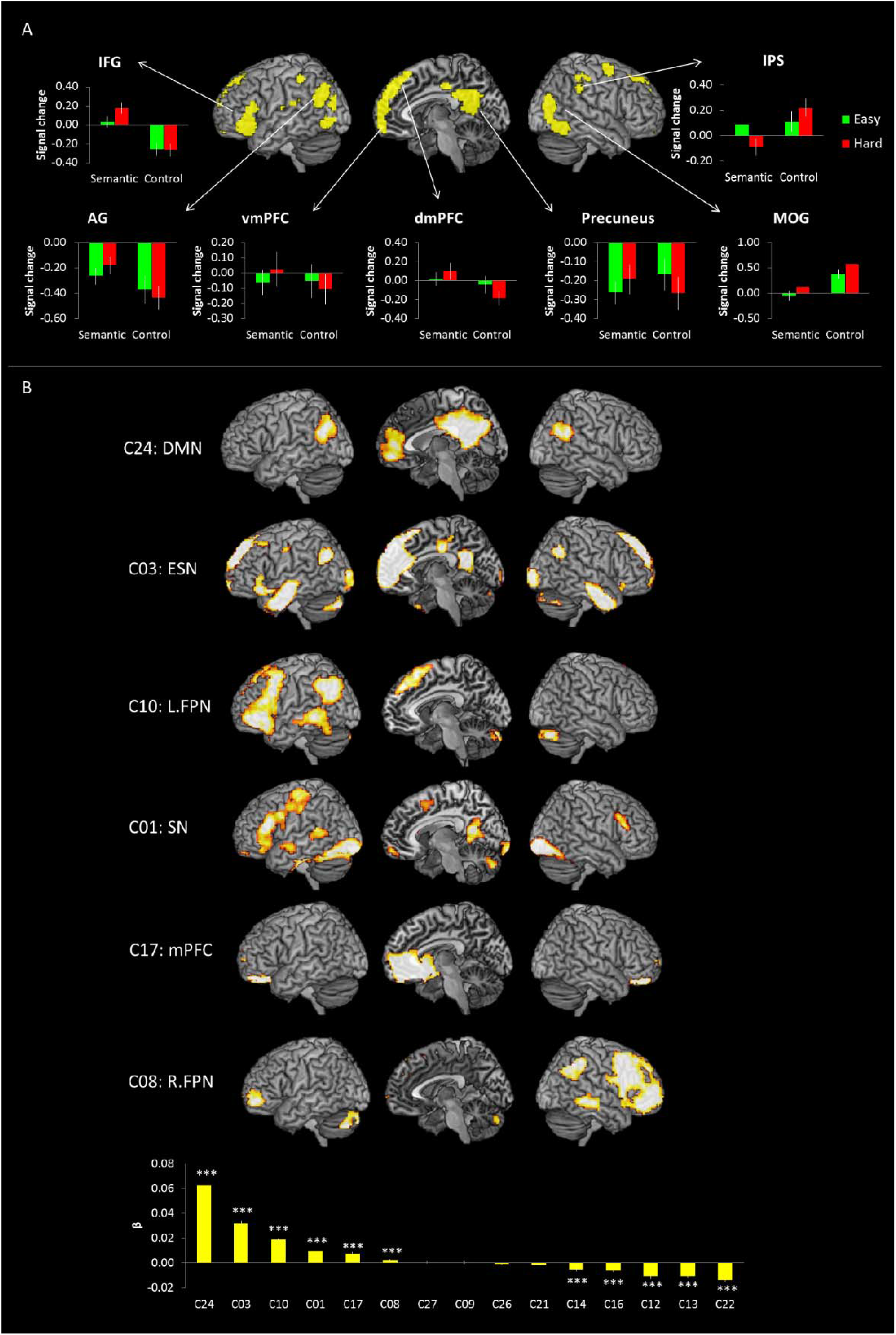
A) The GLM results of the interaction between task and difficulty. Green bar represent regional signal changes for the easy condition and red bar for the hard condition. B) The spatial regression results. The spatial regression analysis identified that C01, C03, C08, C10, C17 and C24 were significantly associated with the brain activation map of the interaction between task and difficulty. *** p _FDR- corrected_ < 0.001

In order to evaluate which ICs is overlapping with the brain map of interaction, we performed the spatial regression analysis. The results revealed that 6 ICs were significantly associated with the GLM results of the effect of interaction (negative value means dissimilarity between the spatial maps so we did not reported, one-sample t-test, p _FDR-corrected_ < 0.001) (Fig 5B Bottom). C24 (DMN), C03 (ESN), C10 (L.FPN), C01 (SN), and C17 (mPFC) overlaps with the regions showed differential activation for semantic processing modulated by task difficulty. These networks were involved in semantic processing in previous analyses (Fig 2 & 3). C08 (R.FPN) showed significantly similarity with the regions responded for control visual processing modulated by its difficulty (Fig 2 & Table 2).

### Functional network connectivity (FNC)

We examined the internetwork connectivity between ICs showed a significant involvement in the interaction of task and difficulty (Fig 6). Correlation analysis was performed between the time-course of the 6 ICs (SN, ESN, R.FPN, L.FPN, mPFC, and DMN). C01 (SN) was significantly correlated with C03 (ESN) and C10 (L.FPN), whereas anti-correlated with C24 (DMN). C03 (ESN) was decoupled with C08 (R.FPN) and coupled with C10 (L.FPN), C17 (mPFC), and C24 (DMN). C08 (R.FPN) was significantly related with C10 (L.FPN). C10 (L.FPN) was positively correlated with C24 (DMN). The results are displayed in Fig 6A.

**Figure 6.**
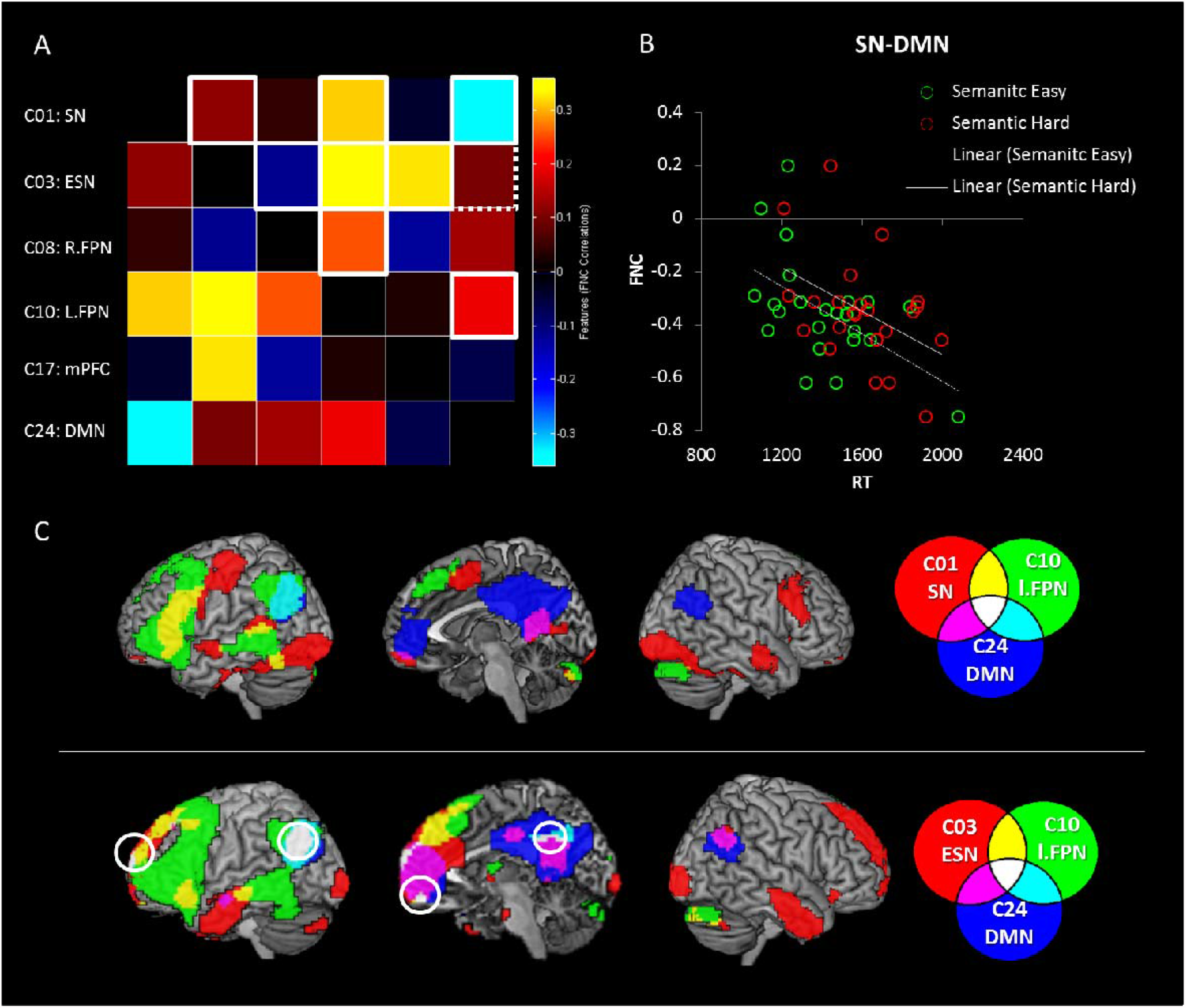
A) The results of FNC between the networks involved in task and difficulty. Red colours indicate positive coupling and Blue colours indicate negative coupling. White box represents a significant FNC between networks (p _FDR-corrected_ < 0.05). Dotted white box represents a marginally significant FNC (p _FDR-corrected_ = 0.07). B) The relationship between the FNC (SN-DMN) and semantic task performance. Green and red circles represent individual performance during easy and hard semantic conditions, respectively. C) The overlapping between domain-specific (semantic processing) and domain-general networks.

To explore the relationship between FNC and task performance, we conducted a correlation analysis. We found that the internetwork connectivity between the SN and DMN was significantly correlated with semantic task performance (Fig 6B). Individuals with less decoupling between them performed the semantic task better (faster RT) regardless of task difficulty (easy: r = -0.54, p _FDR- corrected_ < 0.01; hard: r = -0.44, p _FDR-corrected_ < 0.05).

### Overlapping networks

The ICA decomposes the fMRI data into multiple components with independent sources of variance. Although these components are independent, they can overlap partially. Figure 6C shows the overlapping regions between domain-specific (semantic processing) and domain-general networks. The SN overlapped with the L.FPN in the left IFG and pMTG and with the DMN in the mPFC and PCC. The left AG was common for the L.FPN and the DMN. The ESN overlapped with the L.PFN in the ventral IFG, dorsomedial PFC and MTG and with the DMN in the mPFC, PCC, and the right AG. The left AG, vmPFC, dmPFC and PCC were common regions for all three networks.

## Discussion

The aim of this study was to investigate the neural dynamics of functional brain networks supporting human higher cognition - semantic cognition. ICA revealed that semantic cognition requires the coordination of multiple brain networks subserving cognitive processes. Our task difficulty manipulation segregated and integrated domain-specific (semantic networks) and domain-general systems including the FPN and DMN. The results revealed that semantic system was modulated by difficulty manipulation, demonstrating two independent networks (SN and ESN) with differential contribution to easy and hard semantic processing. The DMN was deactivated during semantic processing but a subset of the DMN, mPFC was activated only for demanding semantic processing. These domain-specific and domain-general networks cooperated with each other during semantic processing. We found that the internetwork connectivity between the SN and DMN is crucial for effective semantic cognition. These findings suggest that semantic cognition is founded on a flexible, dynamic system, revealing its resilience against task demands (Jung and Lambon Ralph 2016; Jung *et al*. 2021). Our results reveal large-scale neural dynamics in the brain such that human higher cognition is achieved by the integration of multiple neural networks, which can be segregated into several sub-networks, serving distinct cognitive functions depending on task demands.

The semantic network is a widely distributed network encompassing the frontal, temporal and parietal cortices, by reflecting its multi-faceted nature of semantic cognition (Binder *et al*. 2009; Lambon Ralph *et al*. 2017). Our ICA revealed two distinctive semantic networks: the left-lateralized semantic network (SN: C01) and the extended semantic network (ESN: C03). The left-lateralized semantic network was activated for the semantic tasks and more engaged in the easy semantic processing. It includes the key semantic regions in the left hemisphere including the ventral ATL – the semantic representation hub and IFG as well as pMTG – semantic control regions (Jefferies and Lambon Ralph 2006; Noonan *et al*. 2013; Lambon Ralph *et al*. 2017). The ESN also was positively engaged in both semantic conditions but preferred to the hard semantic condition. The ESN is a bilateral system including the bilateral ventrolateral ATL, ventral IFG, AG, mPFC, cingulate cortex, and precuneus. This bilateral semantic system has been observed in previous studies (Humphreys *et al*. 2015; Rice, Lambon Ralph, et al. 2015; Jung and Lambon Ralph 2016; Humphreys and Lambon Ralph 2017; Jung *et al*. 2021). Our fMRI combined with repetitive transcranial magnetic stimulation studies demonstrated that the disturbance in the semantic system from the left ATL stimulation induced the up-regulation in the right ATL (Binney and Lambon Ralph 2014) and increased ATL-connectivity (Jung and Lambon Ralph 2016). Our recent fMRI investigation with task difficulty manipulation revealed the bilateral ATL activation as well as increased ATL-interhemispheric connectivity during demanding semantic processing (Jung *et al*. 2021). In the hard semantic trials, the target (e.g., grape) and the foil (e.g., bluebell) were from a same or similar category so there was increased competition between them to relate to the probe (e.g., carrot). For effective semantic processing, it requires the increased involvement of the semantic hub to have a fine appreciation of shades of meaning, which leads to increased regional activity in the both ATLs and increased ATL interhemispheric-connectivity. Importantly, patients with semantic dementia with bilateral ATL atrophy showed severe semantic impairments, whereas patients with unilateral ATL damage showed relatively preserved semantic function (Lambon Ralph, Cipolotti, et al. 2010; Ding et al. 2020). These studies suggest that the bilateral ATL hubs is crucial to sustain semantic performance, making the system robust to damage or task demands (Warren et al. 2009; Jung and Lambon Ralph 2016; Jung *et al*. 2021). Here, our ICA results support the notion that semantic representation is founded on an intrinsically bilateral ATL system (Lambon Ralph *et al*. 2017).

The ESN overlaps with two domain-general networks: L.FPN and DMN (Fig 6C). The L.FPN is a task-active network involved in executive control (Duncan 2010; Zanto and Gazzaley 2013). The ESN overlaps with the L.FPN in the left ventral IFG, dmPFC, MTG, and AG. The ventral IFG and pMTG showed increased activation during demanding semantic tasks (Noonan *et al*. 2013; Jung *et al*. 2021), whereas the DLPFC and IPS were activated for demanding conditions regardless of tasks (Badre *et al*. 2005; Nagel *et al*. 2008). Our ROI analysis demonstrated that the left ventral IFG and pMTG showed activation during semantic processing (more activation for the hard semantic condition) but deactivation during control processing. The overlapping ventral IFG and pMTG within the ESN are associated with the regulation of semantic representation as semantic control regions (Lambon Ralph *et al*. 2017). In addition, the ESN overlaps with the DMN in the bilateral AG, mPFC, and PCC/precuneus. These regions were deactivated during goal-directed tasks (Greicius et al. 2003) but were activated for semantic processing (Binder *et al*. 1999; Binder *et al*. 2009). Specifically, the subsets of DMN showed less deactivation during semantic processing compared to non-semantic processing (Wirth *et al*. 2011; Jung *et al*. 2021) but the PCC and AG was deactivated for all tasks with the level of deactivation related to task difficulty (Humphreys *et al*. 2015; Humphreys and Lambon Ralph 2017). Our ROI analysis (Fig 5A) revealed that these DMN regions showed deactivation for semantic and control tasks but less deactivation (in the AG and PCC/precuneus) or even activation (mPFC) for demanding semantic processing. Furthermore, our ICA analysis extracted the mPFC (C17) as an independent network from the DMN. This network was activated only for demanding semantic processing and positively correlated with the ESN. These overlapping regions between the ESN, L.FPN, and DMN were found in the GLM analysis of the interaction between task and difficulty (Fig 5A). Our results demonstrated that these regions were sensitive to both task and difficulty and ICA unveiled a distinct network composed of these regions – ESN, a hybrid network embracing semantic and domain-general processing. These findings supports the FPN and DMN are a dynamic neural system consisting of several subsystems, serving distinct cognitive functions (Buckner *et al*. 2008; Laird et al. 2009; Andrews-Hanna et al. 2010; Camilleri et al. 2018).

Our results might have significant implications for interpretation of the relationship between the SN and DMN. Due to the uncontrolled rich conceptual processing at rest, there is extensive overlapping between the SN and DMN (Binder *et al*. 1999; Buckner *et al*. 2008; Binder *et al*. 2009). Recent fMRI studies with controlled tasks and comparison of task and rest data have demonstrated the sub-networks of the DMN in relation to various cognitive processing (Andrews-Hanna *et al*. 2010; Wirth *et al*. 2011; Hyatt et al. 2015). For example, Hyatt and colleagues (2015) employed a social and semantic task to decompose the DMN. Using ICA, they found 10 sub-networks within the DMN. They reported that the dmPFC and PCC sub-networks were specifically modulated by semantic memory recall. Similarly, our data showed that ESN, a hybrid network, was activated for semantic processing with different level of involvement depending on task difficulty and a sub-network of the DMN (C17: mPFC) was recruited for the hard semantic processing. These results seem to suggest the functional integration between the SN and DMN for semantic processing. However, we found that the SN (C01) was decoupled with DMN during semantic processing (Fig 6A). Furthermore, the degree of decoupling was correlated with semantic task performance. Individuals with less decoupling between them performed better semantic task better. A potential explanation may be that, although demanding semantic processing fractionises the DMN, a subset or a region of it participates in the processing, and expands the task-related networks. At the same time, the core SN might synchronise itself more for efficient semantic processing, by releasing the suppression toward the task-negative system. Our findings suggest the anti-correlated relationship between the core task-specific and task-negative networks still remains and tends to reduce its decoupling for demanding processing.

ICA approach decomposed the FPN into two networks, showing differential modulation according tasks (semantic vs. non-semantic): L.FPN and R.FPN. The FPN is a multiple-demand (MD) system showing conjoint activation across multiple demands (Duncan 2010). It includes the pre-SMA, ACC/MCC, insular, IPS and inferior frontal sulcus (IFS) bilaterally, contributing to executive processing across tasks (Fedorenko et al. 2013). However, our data demonstrated that this system was divided into two subsets depending on a task. The L.FPN was activated during semantic processing but deactivated during visuo-spatial processing, whereas R.FPN showed the opposite pattern of task involvement. Furthermore, L.FPN was synchronised with the SN, ESN, R.FPN and DMN. In contrary, R.FPN was de-synchronised with the ESN. Possibly, our language-based semantic task induced more involvement of the left-lateralized FPN and the rest of system was allocated for domain-general processing (non-semantic). Our findings support the recent view of domain-general systems that it can be reorganised depending on specific demands by fractionising itself or recruiting additional regions (Camilleri *et al*. 2018).

We found the PSN (C22) was positively engaged in semantic processing. The PSN is a bilateral system including the superior temporal gyrus, insular, RO, and SMG. This network has been implicated as a language comprehension system, especially for phonological/speech processing (Hickok and Poeppel 2007). It has been reported that both phonological and semantic system were recruited for successful visual word recognition (Price 2012; Pattamadilok et al. 2017). Therefore, this network can participate in semantic processing as a key part of language comprehension.

In addition to cortical networks, the thalamus network (C27) also participated in semantic processing, in particular, for the demanding condition. This network includes thalamus, putamen and pallidum. This system has been considered as a station relaying all incoming information from outside world to the cortex (McCormick and Bal 1994). Several studies suggested both the thalamus and the putamen are a part of the MD system that links different regions via cortico-striatal-thalamic circuits (Alexander et al. 1986; Bonelli and Cummings 2007; Camilleri *et al*. 2018). Evidence for cortico-thalamic language processing has also been provided by a study with simultaneous depth and scalp recordings in the context of deep brain stimulation (Wahl et al. 2008). During syntactic processing, language-related potentials (LRP) were identified in the ventrolateral thalamus. During semantic processing, both cortical and thalamic LRPs appeared. Taken together, our data suggests that the thalamus system may be recruited for semantic processing in parallel to domain-general processing, operated through cortico-striatal-thalamic circuits (Ullman 2006).

## Conflict of interest

The authors declare no competing financial interests.

## Acknowledgments

This research was supported by a Beacon Anne McLaren Research Fellowship (University of Nottingham) to JJ and an Advanced ERC award (GAP: 670428 - BRAIN2MIND_NEUROCOMP), MRC programme grant (MR/R023883/1), and intramural funding (MC_UU_00005/18) to MALR.

## Notes

### Competing Interest Statement

The authors have declared no competing interest.

## References

Alexander GE, DeLong MR, Strick PL. 1986. Parallel organization of functionally segregated circuits linking basal ganglia and cortex. Annu Rev Neurosci. 9:357–381.

Andrews-Hanna JR, Reidler JS, Sepulcre J, Poulin R, Buckner RL. 2010. Functional-Anatomic Fractionation of the Brain’s Default Network. Neuron. 65:550–562.

Ashburner J. 2007. A fast diffeomorphic image registration algorithm. Neuroimage. 38:95–113.

Badre D, Poldrack RA, Pare-Blagoev EJ, Insler RZ, Wagner AD. 2005. Dissociable controlled retrieval and generalized selection mechanisms in ventrolateral prefrontal cortex. Neuron. 47:907–918.

Badre D, Wagner AD. 2002. Semantic retrieval, mnemonic control, and prefrontal cortex. Behav Cogn Neurosci Rev. 1:206–218.

Bajada CJ, Haroon HA, Azadbakht H, Parker GJM, Lambon Ralph MA, Cloutman LL. 2017. The tract terminations in the temporal lobe: Their location and associated functions. Cortex. 97:277–290.

Bell AJ, Sejnowski TJ. 1995. An Information Maximization Approach to Blind Separation and Blind Deconvolution. Neural Comput. 7:1129–1159.

Binder JR, Conant LL, Humphries CJ, Fernandino L, Simons SB, Aguilar M, Desai RH. 2016. Toward a brain-based componential semantic representation. Cogn Neuropsychol. 33:130–174.

Binder JR, Desai RH, Graves WW, Conant LL. 2009. Where is the semantic system? A critical review and meta-analysis of 120 functional neuroimaging studies. Cereb Cortex. 19:2767–2796.

Binder JR, Frost JA, Hammeke TA, Bellgowan PS, Rao SM, Cox RW. 1999. Conceptual processing during the conscious resting state. A functional MRI study. J Cogn Neurosci. 11:80–95.

Binney RJ, Embleton KV, Jefferies E, Parker GJ, Ralph MA. 2010. The ventral and inferolateral aspects of the anterior temporal lobe are crucial in semantic memory: evidence from a novel direct comparison of distortion-corrected fMRI, rTMS, and semantic dementia. Cereb Cortex. 20:2728–2738.

Binney RJ, Lambon Ralph MA. 2014. Using a combination of fMRI and anterior temporal lobe rTMS to measure intrinsic and induced activation changes across the semantic cognition network. Neuropsychologia.

Binney RJ, Parker GJ, Lambon Ralph MA. 2012. Convergent connectivity and graded specialization in the rostral human temporal lobe as revealed by diffusion-weighted imaging probabilistic tractography. J Cogn Neurosci. 24:1998–2014.

Bonelli RM, Cummings JL. 2007. Frontal-subcortical circuitry and behavior. Dialogues Clin Neurosci. 9:141–151.

Bozeat S, Lambon Ralph MA, Patterson K, Garrard P, Hodges JR. 2000. Non-verbal semantic impairment in semantic dementia. Neuropsychologia. 38:1207–1215.

Bressler SL, Kelso JA. 2001. Cortical coordination dynamics and cognition. Trends Cogn Sci. 5:26–36.

Buckner RL, Andrews-Hanna JR, Schacter DL. 2008. The brain’s default network: anatomy, function, and relevance to disease. Ann N Y Acad Sci. 1124:1–38.

Calhoun VD, Adali T, McGinty VB, Pekar JJ, Watson TD, Pearlson GD. 2001. fMRI activation in a visual-perception task: network of areas detected using the general linear model and independent components analysis. Neuroimage. 14:1080–1088.

Calhoun VD, Adali T, Pearlson GD, Pekar JJ. 2001. A method for making group inferences from functional MRI data using independent component analysis. Hum Brain Mapp. 14:140–151.

Camilleri JA, Muller VI, Fox P, Laird AR, Hoffstaedter F, Kalenscher T, Eickhoff SB. 2018. Definition and characterization of an extended multiple-demand network. Neuroimage. 165:138–147.

Chen Y, Shimotake A, Matsumoto R, Kunieda T, Kikuchi T, Miyamoto S, Fukuyama H, Takahashi R, Ikeda A, Lambon Ralph MA. 2016. The ‘when’ and ‘where’ of semantic coding in the anterior temporal lobe: Temporal representational similarity analysis of electrocorticogram data. Cortex. 79:1–13.

Coccia M, Bartolini M, Luzzi S, Provinciali L, Ralph MA. 2004. Semantic memory is an amodal, dynamic system: Evidence from the interaction of naming and object use in semantic dementia. Cogn Neuropsychol. 21:513–527.

Cohen JR, D’Esposito M. 2016. The Segregation and Integration of Distinct Brain Networks and Their Relationship to Cognition. J Neurosci. 36:12083–12094.

Corbett F, Jefferies E, Ehsan S, Lambon Ralph MA. 2009. Different impairments of semantic cognition in semantic dementia and semantic aphasia: evidence from the non-verbal domain. Brain. 132:2593– 2608.

Ding J, Chen K, Liu H, Huang L, Chen Y, Lv Y, Yang Q, Guo Q, Han Z, Lambon Ralph MA. 2020. A unified neurocognitive model of semantics language social behaviour and face recognition in semantic dementia. Nat Commun. 11:2595.

Domagalik A, Beldzik E, Fafrowicz M, Oginska H, Marek T. 2012. Neural networks related to prosaccades and anti-saccades revealed by independent component analysis. Neuroimage. 62:1325– 1333.

Duncan J. 2010. The multiple-demand (MD) system of the primate brain: mental programs for intelligent behaviour. Trends Cogn Sci. 14:172–179.

Fedorenko E, Duncan J, Kanwisher N. 2012. Language-selective and domain-general regions lie side by side within Broca’s area. Curr Biol. 22:2059–2062.

Fedorenko E, Duncan J, Kanwisher N. 2013. Broad domain generality in focal regions of frontal and parietal cortex. Proc Natl Acad Sci U S A. 110:16616–16621.

Geranmayeh F, Wise RJ, Mehta A, Leech R. 2014. Overlapping networks engaged during spoken language production and its cognitive control. J Neurosci. 34:8728–8740.

Greicius MD, Krasnow B, Reiss AL, Menon V. 2003. Functional connectivity in the resting brain: A network analysis of the default mode hypothesis. P Natl Acad Sci USA. 100:253–258.

Halai AD, Welbourne SR, Embleton K, Parkes LM. 2014. A comparison of dual gradient-echo and spin-echo fMRI of the inferior temporal lobe. Hum Brain Mapp. 35:4118–4128.

Hickok G, Poeppel D. 2007. The cortical organization of speech processing. Nat Rev Neurosci. 8:393– 402.

Humphreys GF, Hoffman P, Visser M, Binney RJ, Lambon Ralph MA. 2015. Establishing task-and modality-dependent dissociations between the semantic and default mode networks. Proc Natl Acad Sci U S A. 112:7857–7862.

Humphreys GF, Lambon Ralph MA. 2017. Mapping Domain-Selective and Counterpointed Domain-General Higher Cognitive Functions in the Lateral Parietal Cortex: Evidence from fMRI Comparisons of Difficulty-Varying Semantic Versus Visuo-Spatial Tasks, and Functional Connectivity Analyses. Cereb Cortex. 27:4199–4212.

Hyatt CJ, Calhoun VD, Pearlson GD, Assaf M. 2015. Specific default mode subnetworks support mentalizing as revealed through opposing network recruitment by social and semantic FMRI tasks. Hum Brain Mapp. 36:3047–3063.

Jackson RL, Hoffman P, Pobric G, Lambon Ralph MA. 2016. The Semantic Network at Work and Rest: Differential Connectivity of Anterior Temporal Lobe Subregions. J Neurosci. 36:1490–1501.

Jackson RL, Hoffman P, Pobric G, Ralph MAL. 2015. The Nature and Neural Correlates of Semantic Association versus Conceptual Similarity. Cerebral Cortex. 25:4319–4333.

Jafri MJ, Pearlson GD, Stevens M, Calhoun VD. 2008. A method for functional network connectivity among spatially independent resting-state components in schizophrenia. Neuroimage. 39:1666–1681.

Jefferies E, Lambon Ralph MA. 2006. Semantic impairment in stroke aphasia versus semantic dementia: a case-series comparison. Brain. 129:2132–2147.

Jung J, Cloutman LL, Binney RJ, Lambon Ralph MA. 2017. The structural connectivity of higher order association cortices reflects human functional brain networks. Cortex.

Jung J, Lambon Ralph MA. 2016. Mapping the Dynamic Network Interactions Underpinning Cognition: A cTBS-fMRI Study of the Flexible Adaptive Neural System for Semantics. Cereb Cortex. 26:3580– 3590.

Jung JY, Rice GE, Lambon Ralph MA. 2021. The neural bases of resilient semantic system: evidence of variable neuro-displacement in cognitive systems. Brain Struct Funct. 226:1585–1599.

Kim DI, Manoach DS, Mathalon DH, Turner JA, Mannell M, Brown GG, Ford JM, Gollub RL, White T, Wible C, Belger A, Bockholt HJ, Clark VP, Lauriello J, O’Leary D, Mueller BA, Lim KO, Andreasen N, Potkin SG, Calhoun VD. 2009. Dysregulation of working memory and default-mode networks in schizophrenia using independent component analysis, an fBIRN and MCIC study. Hum Brain Mapp. 30:3795–3811.

Laird AR, Eickhoff SB, Li K, Robin DA, Glahn DC, Fox PT. 2009. Investigating the functional heterogeneity of the default mode network using coordinate-based meta-analytic modeling. J Neurosci. 29:14496–14505.

Lambon Ralph MA, Cipolotti L, Manes F, Patterson K. 2010. Taking both sides: do unilateral anterior temporal lobe lesions disrupt semantic memory? Brain. 133:3243–3255.

Lambon Ralph MA, Jefferies E, Patterson K, Rogers TT. 2017. The neural and computational bases of semantic cognition. Nat Rev Neurosci. 18:42–55.

Lambon Ralph MA, Sage K, Jones RW, Mayberry EJ. 2010. Coherent concepts are computed in the anterior temporal lobes. P Natl Acad Sci USA. 107:2717–2722.

Martin A. 2016. GRAPES-Grounding representations in action, perception, and emotion systems: How object properties and categories are represented in the human brain. Psychon Bull Rev. 23:979– 990.

McCormick DA, Bal T. 1994. Sensory gating mechanisms of the thalamus. Curr Opin Neurobiol. 4:550–556.

McKeown MJ, Sejnowski TJ. 1998. Independent component analysis of fMRI data: examining the assumptions. Hum Brain Mapp. 6:368–372.

Nagel IE, Schumacher EH, Goebel R, D’Esposito M. 2008. Functional MRI investigation of verbal selection mechanisms in lateral prefrontal cortex. Neuroimage. 43:801–807.

Noonan KA, Jefferies E, Visser M, Lambon Ralph MA. 2013. Going beyond inferior prefrontal involvement in semantic control: evidence for the additional contribution of dorsal angular gyrus and posterior middle temporal cortex. J Cogn Neurosci. 25:1824–1850.

Oldfield RC. 1971. The assessment and analysis of handedness: the Edinburgh inventory. Neuropsychologia. 9:97–113.

Pascual B, Masdeu JC, Hollenbeck M, Makris N, Insausti R, Ding SL, Dickerson BC. 2015. Large-scale brain networks of the human left temporal pole: a functional connectivity MRI study. Cereb Cortex. 25:680–702.

Pattamadilok C, Chanoine V, Pallier C, Anton JL, Nazarian B, Belin P, Ziegler JC. 2017. Automaticity of phonological and semantic processing during visual word recognition. Neuroimage. 149:244–255.

Patterson K, Nestor PJ, Rogers TT. 2007. Where do you know what you know? The representation of semantic knowledge in the human brain. Nat Rev Neurosci. 8:976–987.

Peelen MV, Caramazza A. 2012. Conceptual object representations in human anterior temporal cortex. J Neurosci. 32:15728–15736.

Price CJ. 2012. A review and synthesis of the first 20 years of PET and fMRI studies of heard speech, spoken language and reading. Neuroimage. 62:816–847.

Raichle ME, MacLeod AM, Snyder AZ, Powers WJ, Gusnard DA, Shulman GL. 2001. A default mode of brain function. Proc Natl Acad Sci U S A. 98:676–682.

Rice GE, Hoffman P, Lambon Ralph MA. 2015. Graded specialization within and between the anterior temporal lobes. Ann N Y Acad Sci.

Rice GE, Lambon Ralph MA, Hoffman P. 2015. The Roles of Left Versus Right Anterior Temporal Lobes in Conceptual Knowledge: An ALE Meta-analysis of 97 Functional Neuroimaging Studies. Cereb Cortex. 25:4374–4391.

Shine JM, Bissett PG, Bell PT, Koyejo O, Balsters JH, Gorgolewski KJ, Moodie CA, Poldrack RA. 2016. The Dynamics of Functional Brain Networks: Integrated Network States during Cognitive Task Performance. Neuron. 92:544–554.

Thompson-Schill SL, D’Esposito M, Aguirre GK, Farah MJ. 1997. Role of left inferior prefrontal cortex in retrieval of semantic knowledge: a reevaluation. Proc Natl Acad Sci U S A. 94:14792–14797.

Uddin LQ. 2017. Salience Network of the Human Brain. Salience Network of the Human Brain.1-34.

Ullman MT. 2006. Is Broca’s area part of a basal ganglia thalamocortical circuit? Cortex. 42:480–485.

Vossel S, Geng JJ, Fink GR. 2014. Dorsal and Ventral Attention Systems: Distinct Neural Circuits but Collaborative Roles. Neuroscientist. 20:150–159.

Wahl M, Marzinzik F, Friederici AD, Hahne A, Kupsch A, Schneider GH, Saddy D, Curio G, Klostermann F. 2008. The human thalamus processes syntactic and semantic language violations. Neuron. 59:695– 707.

Warren JE, Crinion JT, Lambon Ralph MA, Wise RJ. 2009. Anterior temporal lobe connectivity correlates with functional outcome after aphasic stroke. Brain. 132:3428–3442.

Whitney C, Kirk M, O’Sullivan J, Lambon Ralph MA, Jefferies E. 2011. The neural organization of semantic control: TMS evidence for a distributed network in left inferior frontal and posterior middle temporal gyrus. Cereb Cortex. 21:1066–1075.

Wirth M, Jann K, Dierks T, Federspiel A, Wiest R, Horn H. 2011. Semantic memory involvement in the default mode network: a functional neuroimaging study using independent component analysis. Neuroimage. 54:3057–3066.

Zanto TP, Gazzaley A. 2013. Fronto-parietal network: flexible hub of cognitive control. Trends Cogn Sci. 17:602–603.

